# Using virtual reality and thermal imagery to improve statistical modelling of vulnerable and protected species

**DOI:** 10.1101/645291

**Authors:** Catherine Leigh, Grace Heron, Ella Wilson, Taylor Gregory, Samuel Clifford, Jacinta Holloway, Miles McBain, Felipé Gonzalez, James McGree, Ross Brown, Kerrie Mengersen, Erin E. Peterson

## Abstract

Biodiversity loss and sparse observational data mean that critical conservation decisions may be based on little to no information. Emerging technologies, such as airborne thermal imaging and virtual reality, may facilitate species monitoring and improve predictions of species distribution. Here we combined these two technologies to predict the distribution of koalas, specialized arboreal foliovores facing population declines in many parts of eastern Australia. For a study area in southeast Australia, we complemented ground-survey records with presence and absence observations from thermal-imagery obtained using Remotely-Piloted Aircraft Systems. These field observations were further complemented with information elicited from koala experts, who were immersed in 360-degree images of the study area. The experts were asked to state the probability of observing a koala at sites they viewed and to assign each probability a confidence rating. We fit logistic regression models to the ground survey data and the ground plus thermal-imagery survey data and a beta regression model to the expert elicitation data. We then combined parameter estimates from the expert-elicitation model with those from each of the survey models to predict koala presence and absence in the study area. The model that combined the ground, thermal-imagery and expert-elicitation data substantially reduced the uncertainty around parameter estimates and increased the accuracy of classifications (koala presence vs absence), relative to the model based on ground-survey data alone. Our findings suggest that data elicited from experts using virtual reality technology can be combined with data from other emerging technologies, such as airborne thermal-imagery, using traditional statistical models, to increase the information available for species distribution modelling and the conservation of vulnerable and protected species.

## Introduction

In the face of unprecedented biodiversity loss, critical decisions are needed on the conservation of vulnerable and protected species [1,2]. Unfortunately, information is seldom available or dense enough in space and time to effectively inform those decisions [3]. Monitoring programs often rely on observational records of species collected during ground surveys. However, ground-based detection of vulnerable and protected species is difficult, particularly when species are rare or elusive, and information may be biased towards single or few individuals (e.g. radio-collared animals) or sites with high abundance [4,5]. Furthermore, monitoring large areas using traditional ground-survey methods is logistically and financially infeasible, and while professional monitoring may be time consuming and expensive, volunteer data may be biased and of questionable quality [6,7]. These issues all contribute towards the sparse data problem.

Emerging technologies provide an opportunity to increase the spatial and temporal coverage of data, increase the quality of information gained, potentially lower the cost of sampling, and thereby benefit conservation efforts for species and their habitats. For instance, implementation of top-down thermal imaging captured by Remotely-Piloted Aircraft Systems (RPAS), commonly known as drones or unpiloted aerial vehicles, can provide a cost-effective alternative to species counting [8,9]. Alternatively, virtual reality (VR) can create immersive experiences of field conditions used to gather expert information both cost-effectively and with relative ease [10]. By bringing the environment to the wearer, the immersive experiences are expected to improve elicitation responses due to the priming of associated visual memories laid down while physically present in similar environments [11,12]. The resulting information can then be incorporated into quantitative analyses as data or informative priors (e.g. [13]).

The koala (*Phascolarctos cinereus*) is an iconic native Australian marsupial listed as vulnerable in Queensland, New South Wales and the Australian Capital Territory under the Australian Commonwealth Environmental Protection and Biodiversity Conservation (EPBC) Act 1999 [14,15]. Populations are threatened by multiple and likely interacting factors, including habitat loss and fragmentation, drought, disease, dog attack and vehicle collision [15,16]. Furthermore, these highly specialized, arboreal foliovores are restricted to regions dominated by species within the *Eucalyptus*, *Angophora*, *Corymbia*, *Lophostemon* and/or *Melaleuca* genera, their food-tree species. Consequently, the presence and quality of the food-tree species along with soil quality, which can affect where their preferred trees grow, have been identified as additional factors affecting koala presence [15,16].

Using the koala as a case study, we aimed to demonstrate how thermal-imagery and expert opinions can be harnessed to add value to models based on ground-survey data alone. Few studies have considered combining statistical models with thermal imagery [17] or VR-elicited expert information [18,19]. To our knowledge, this is the first study to use both methods for species distribution modelling in a conservation context. We modelled koala presence and absence based on a suite of habitat characteristics including the presence of food-tree species and distance to potential sources of disturbance (e.g. walking tracks where dogs may be present), and information about habitat suitability elicited from experts immersed in 360-degree imagery. This allowed us to examine whether combining information gained from emerging technologies with ground-survey data would improve (i) understanding of the drivers influencing species presence or absence, and (ii) the accuracy and/or precision of predictions at unsampled locations.

## Materials and methods

### Study area

Alexander Clarke Park in Loganholme, southeast Queensland, Australia (Fig 1) provides habitat for a small population of koalas. The park is approximately 0.20 km^2^, open to the public and contains a fenced-off area for off-leash dogs, multiple playgrounds and walking tracks. The roughly triangular-shaped park is bordered on two sides by the Logan River, and on its third, northwestern side by residential housing (Fig 1). Vegetation in the parkland ranges from dense forest to open scrubland and grass fields.

**Fig 1.**
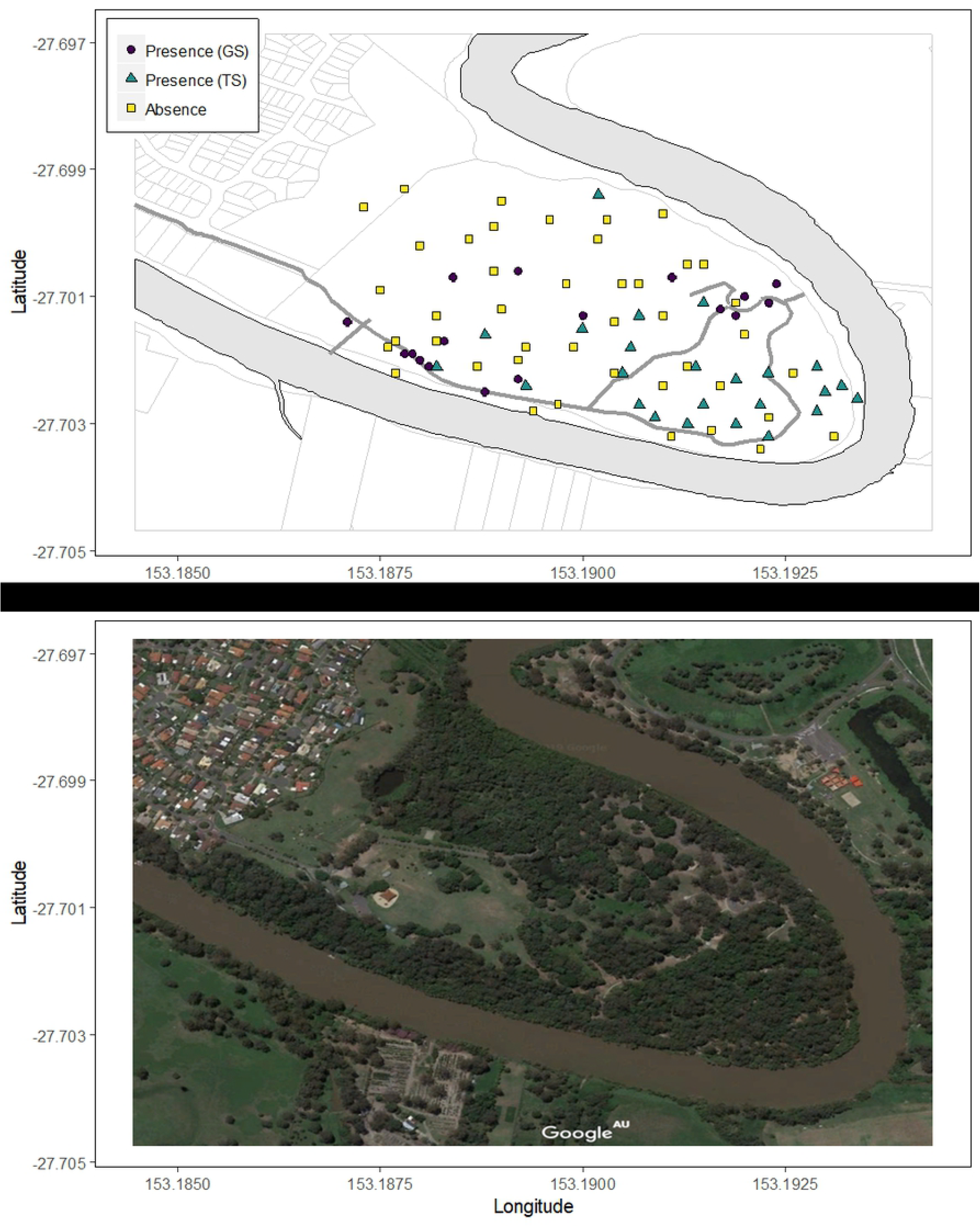
Study area. Alexander Clarke Park in southeast Queensland, Australia. Upper plot: koala presences identified from ground surveys (GS, closed circles) or aerial thermal-imagery surveys (TS, closed triangles) and koala absences (closed squares). Grey shading shows the Logan River, thick grey lines show paths through the park, thin grey lines show Logan City Council lot boundaries, including residential lots in the northwest corner. Lower plot: image data © Google 2019.

### Koala observation data

The complete dataset of 82 koala observations contained 41 unique presences and 41 unique absences (Fig 1; S1 Data). Seventeen of the unique presences were identified from sightings made during ground surveys conducted by citizen scientists between 2012 and 2017 [20] and the project team in December 2017. The remaining 24 unique presences were identified from aerial thermal-imagery surveys conducted in October 2016 and December 2017. On both occasions, thermal imagery was collected during RPAS (DJI M600) flights conducted over the study area for later identification of thermal hotspots as ‘potential’ koalas (Fig 2). A Tau-2 640 captured forward-looking infrared radiometer footage while a Mobius Action Camera and Sony NEX5 captured red-green-blue footage. The RPAS was flown at 60-70 m above ground to accommodate the size of the site within time constraints in the morning (between 5:50 am and 9:20 am; 2016 and 2017) and evening (between 3:53 pm and 5:56 pm; 2016 only). Volunteers (in October 2016) or members of our project team (in December 2017) conducted ground surveys while the footage was captured, noting the geographical coordinates of any koala sightings. Resultant data were then input into a koala detection algorithm [8] to identify koalas and assign confidence ratings between 0 and 1. Eleven koala presences were identified in 2016, and 13 in 2017. These 24 presences had confidence ratings of 0.50 (uncertain hotspot, unconfirmed on the ground), 0.90 (high certainty hotspot, unconfirmed on the ground) or 1.00 (high certainty hotspot, confirmed on the ground; S1 Data). Using all of the survey data collected in December 2017, the 41 unique koala absences were then randomly generated from sites in the study area where koalas were known to be absent.

**Fig 2.**
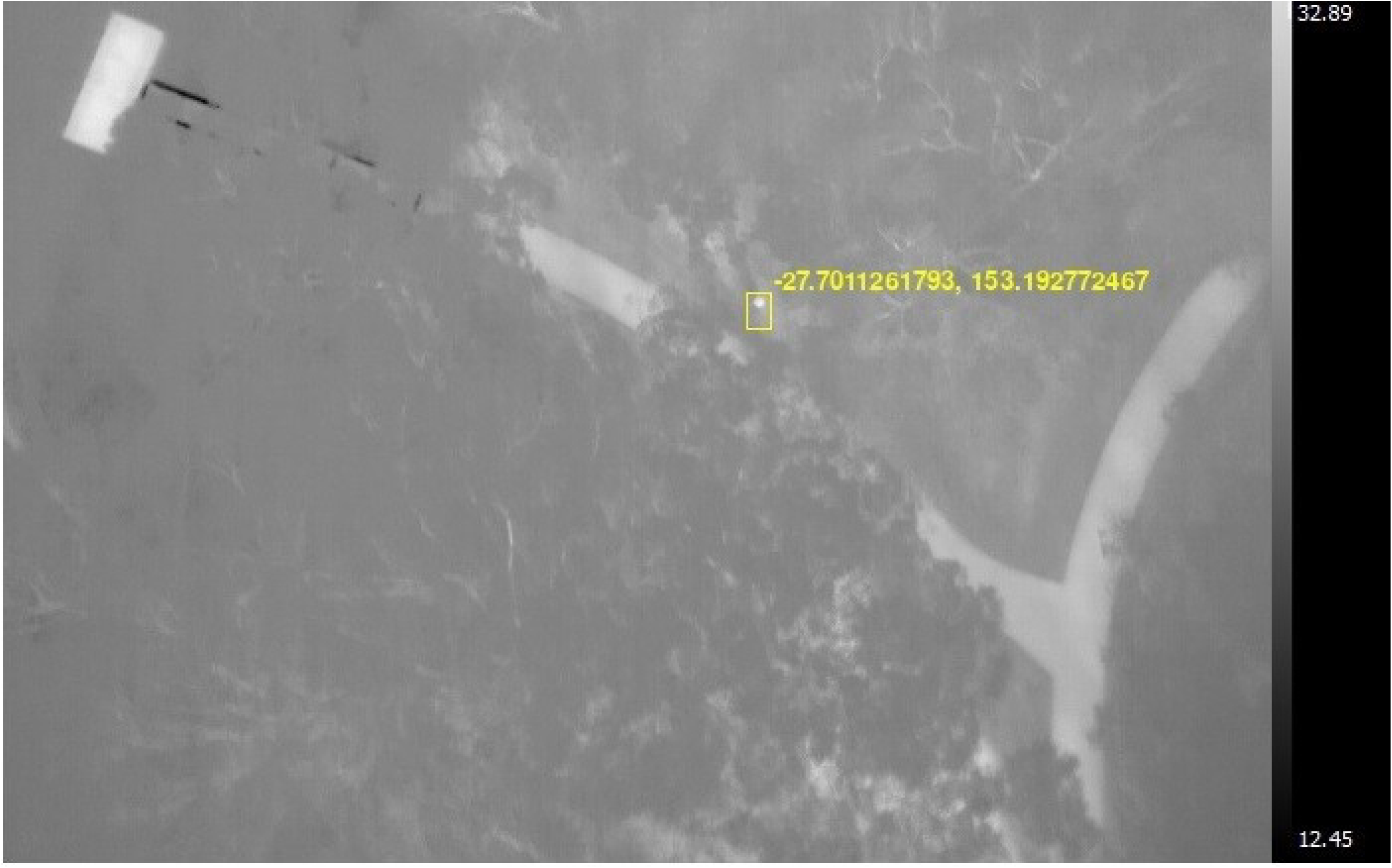
Airborne Thermal Image. Example of a thermal image of a koala captured by the Remotely-Piloted Aircraft Systems.

### Habitat data

Habitat data were collected during field surveys by the project team or derived from freely available geographic information system (GIS) datasets. In December 2017, we estimated the height and density of the tallest vegetation layer in the field, following [21], at each of the 82 koala observation (presence and absence) sites. Height was estimated by measuring the angle of elevation to the top of the tallest tree and the distance between the tree and observer, and then recorded on an ordinal scale of 1 to 5, where 1 > 30 m, 2 =10-30 m, 3 < 10 m, 4 = 2-8 m, and 5 = 0-2 m. Density was based on visual estimates of percentage canopy cover and recorded on an ordinal scale from 1 to 4, where 1 = 70-100%, 2 = 30-70%, 3 = 10-30%, and 4 < 10%.

GIS-based covariates were generated for each of the 82 koala observation sites and for each of 636 additional unobserved prediction sites (see *Model prediction visualization*). This included foliage projective cover (FPC, %) [22], which is the percent of ground cover occupied by the vertical projection of foliage. The percentages were based on dry season (May to October) Landsat-5 TM, Landsat-7 ETM+ and Landsat-8 OLI imagery for the period 1988-2013 [22]. We also generated a binary covariate that represented whether a site contained remnant vegetation dominated by *Eucalyptus* food-tree species (1), or otherwise (0) [23]. Finally, we measured the Euclidean distances (m) to the nearest path [24] and the nearest fresh water (i.e. the Logan River [25]) from each site. The GIS-based covariates were created in R statistical software [26] using the sp [27], raster [28] and geosphere [29] packages.

### Expert elicitation data

Subsets of images that captured the range of koala habitat at the 82 koala observation sites were shown to six koala experts. To generate the subsets, we used cluster analysis (complete-linkage unweighted pair grouping method with arithmetic mean and the Gower distance measure; [30]) to group sites based on vegetation height and density, distance to water, FPC, and koala presence or absence. The resultant clusters contained between 7 and 11 sites each. We created subsets of ten sites each by randomly selecting a site from each cluster. Images of these sites were then captured in December 2017 using a Samsung Gear camera (Fig 3). Finally, we converted these images into 360-degree views using the game engine Unity (Unity Technologies, San Francisco, www.unity3d.com).

**Fig 3.**
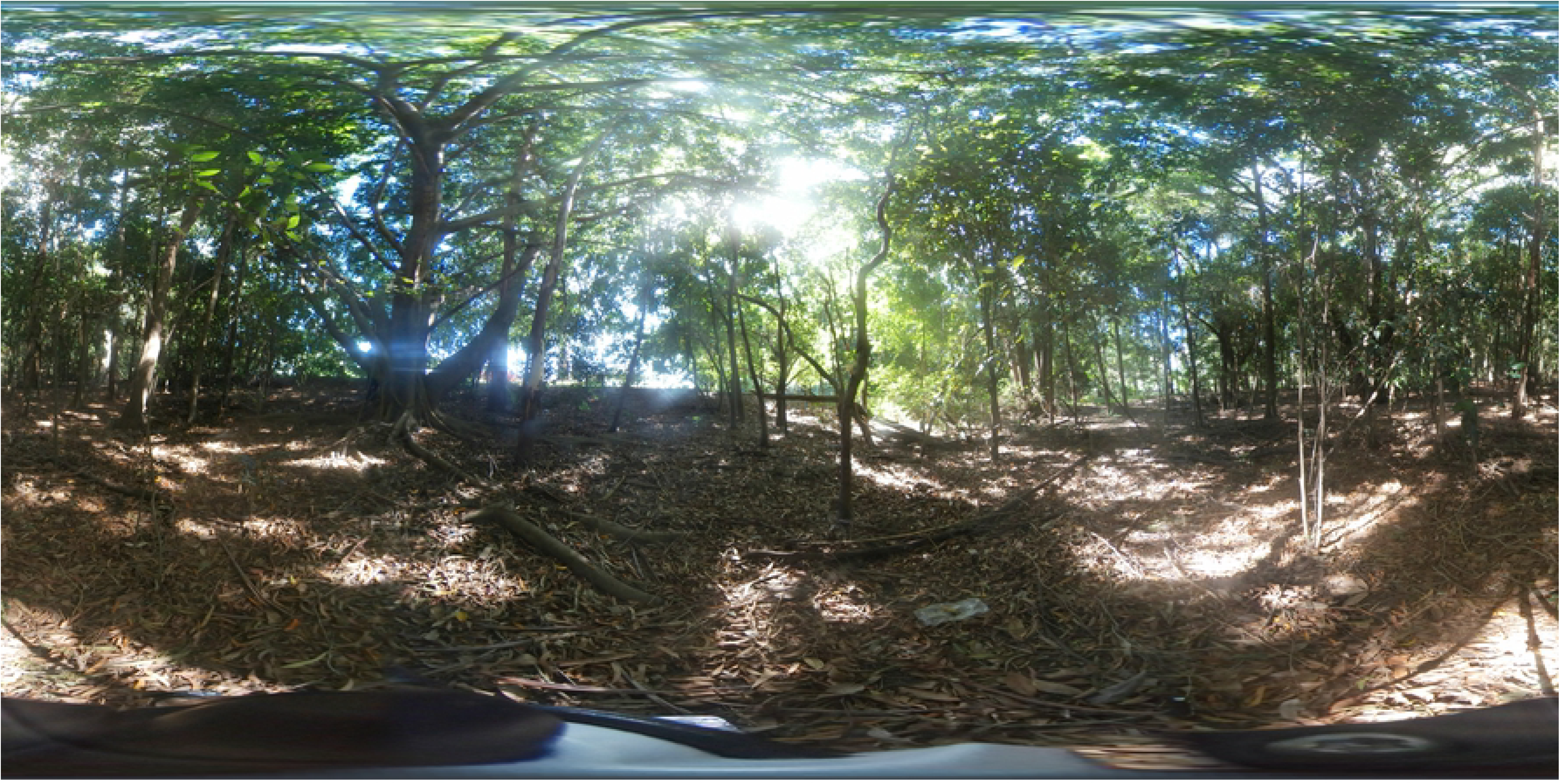
A 360-degree image. Used for virtual reality expert elicitation, showing a site where a koala had been observed during a ground survey. Image by Grace Heron, December 2017.

To elicit information from the experts on koala presence and absence in the study area, we showed each expert ten 360-degree images from a randomly selected subset. The experts included government employees, academic researchers and citizen scientists representing a mix of genders and ranging in age and type of koala expertise (e.g. management, research). Following a standardized protocol (S2 Protocol), we first briefed the experts about the study and provided them with definitions of terms, explanations of probabilities [31], and practice questions while they wore VR-headsets and were immersed in example images. The expert-elicitation interview then followed, in which each expert was asked, for each of the ten images in which they were immersed, (i) “What is the probability that this is a suitable koala habitat?” and (ii) their confidence in each of their answers (very unsure, medium sure or very sure; S2 Protocol). For the purposes of model fitting, the probability an expert gave was assumed to equal the probability of a koala being present at the site being viewed. The same elicitor interviewed each expert, each expert was interviewed separately, and images were shown to each expert in random order.

## Statistical modelling

### Survey-based models

We fit a logistic regression model [32] to the koala observation data from ground and thermal-imagery surveys, using R statistical software [26]. We fit two models: one to the ground-survey data only (the ‘base’ G model, n = 34), and one to the combined ground and thermal-imagery survey data (the GT model, n = 82). We included seven covariates in each model: FPC, remnant *Eucalyptus* vegetation, distance to the nearest path, distance to the nearest fresh water, longitude and latitude. In each case, the covariates were centered and scaled using the means and standard deviations of the respective covariates in the base (G) model. We also included weights in each model based on the confidence in each presence or absence observation: 1.00 for the 17 presences from the ground surveys; 0.50, 0.90 or 1.00 for the 24 presences from the thermal-imagery surveys (as per the confidence ratings from the koala detection algorithm; S1 Data), and 0.90 for the 41 absences. The 0.90 weighting for absences was appropriate because the absences had been generated randomly from sites where koalas had not previously been observed in ground or thermal imagery surveys; as such, there was a small chance that koalas may actually have been present at those sites (i.e. false negatives). The degrees of freedom in the G model were limited because only 34 observations were used to estimate eight parameters (i.e. seven covariates and the intercept). Nevertheless, the model served as a base from which we could compare each of the subsequent models, which included additional data from the thermal-imagery surveys (the GT model), the VR-elicited expert information or both sources (see below).

### Expert elicitation model

We fit a Beta regression model with random effects [33] to the expert elicitation data (the E model, n = 60) using the R package `mgcv` [34]. We matched the elicited answers provided by six experts to the shape parameters of a Beta(*a, b*) model with the probability of observing a koala at a site, 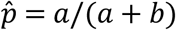, and the total number of koalas expected to be observed, *n*, using the equivalent prior sample method of [35], where 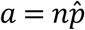 and 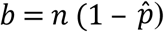 [36]. We used weights of 0.50, 0.75 and 1.00 in the E model. The weights were based on the experts’ confidence in their stated probabilities of not very sure, medium sure, and very sure, respectively (S2 Protocol), and spanned the range of weights used for the survey-based models (i.e. 0.50 to 1.00). We centered and scaled each covariate in the E model (i.e. FPC, remnant *Eucalyptus* vegetation, distance to the nearest path, distance to the nearest fresh water, longitude, latitude) using the means and standard deviations of the respective covariates in the base (G) model. Finally, expert (nominal: one to six) was included in the E model as a random effect.

### Combined models

We combined the estimated parameters and their standard errors from the fitted E model with those from (i) the fitted G model and (ii) the fitted GT model, following [37], to model the distribution of koalas in the study area. This created a combined G_E and a combined GT_E model, respectively. For each of these models, we combined the inverse variance-weighted estimates of the parameter effects [38] from the logistic regression on the observation (*o*) data from the G or GT model, *β*_*o*_, and the beta regression on the elicited (*e*) data from the E model, *β*_*e*_, with the variances 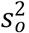, and 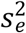, respectively. The combined (*c*) parameter estimates and variances are given by:

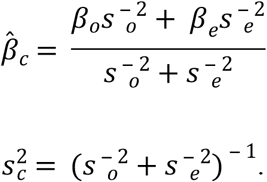

We calculated confidence intervals for the parameters from each of the two combined models (G_E and GT_E) using the respective combined standard error, *s*_*c*_, and the number of degrees of freedom for the *t* statistic being equal to the sum of the residual degrees of freedom from the respective G or GT model and the E model, 𝑣_*c*_ = 𝑣_*o*_ + 𝑣_*e*_, where *α* is the level of statistical significance (0.05):

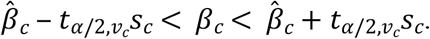

### Predictive performance evaluation

We used a dataset comprising koala observations from the thermal-imagery surveys that had high confidence (weights ≥ 0.90) as a validation dataset to assess the predictive performance of each model (G, GT, G_E and GT_E). As with other datasets, we centered and scaled each of the covariates in this dataset using the means and standard deviations of the respective covariates in the base (G) model. For the G and G_E models, we used the fitted models to make predictions at the validation sites. For the GT and GT_E models, we performed leave-one-out cross validation (LOOCV) whereby a validation site was removed, the GT or GT_E model was fit to the remaining data, and a prediction was then made at the validation site. This process continued until a LOOCV prediction was made at each validation site.

In this modelling scenario, the priority was to identify as many koala presences as possible, rather than to find areas where they were absent. A 0.4 threshold was thus used to classify predictions as either 0 (absence; for prediction values < 0.4), or 1 (presence; for prediction values ≥ 0.4) [39]. We then compared the observations and predictions to evaluate the predictive performance of each model based on classification accuracy (correct predictions divided by the total number of predictions), along with sensitivity (true positives) and specificity (true negatives) and root mean-square prediction error (RMSPE). The larger the classification accuracy, sensitivity and specificity, and smaller the RMSPE, the greater the predictive ability of the model.

### Model prediction visualization

Each of the fitted models (G, GT, G_E and GT_E) was used to make predictions at 636 unobserved sites across the study area. This allowed us to visualize and compare koala distribution across the study area as predicted by each model (G, GT, G_E and GT_E). The covariates in this dataset (i.e. FPC, remnant *Eucalyptus* vegetation, distance to the nearest path, distance to the nearest fresh water, longitude, latitude) were each centered and scaled using the means and standard deviations of the respective covariates in the base (G) model.

## Results

Accuracy of the base G model increased by 75% and RMSPE decreased by 26 % when ground-survey observations were combined with data from the emerging technologies (GT_E model; Table 1). The GT_E model had the greatest accuracy and sensitivity (true presence rate) and smallest RMSPE of all models (Table 1). Although the G model had the greatest specificity (true absence rate), it had exceptionally low sensitivity (0.25), suggesting the model predicted koalas to be absent in most locations. The specificity of the GT_E model was relatively high (at 0.375) given there was no negative impact on the sensitivity of this model (0.937).

**Table 1.** Predictive performance of each model based on accuracy, sensitivity, specificity and root mean-square prediction error (RMSPE). G = ground survey only, GT = ground and thermal-imagery surveys, G_E = combined ground-survey and expert-elicitation, GT_E = combined ground and thermal-imagery surveys and expert-elicitation.

| Model | Accuracy | Sensitivity | Specificity | RMSPE |
| --- | --- | --- | --- | --- |
| G | 0.375 | 0.250 | 0.500 | 0.791 |
| GT | 0.625 | 0.812 | 0.437 | 0.612 |
| G_E | 0.562 | 0.875 | 0.250 | 0.661 |
| GT_E | 0.656 | 0.937 | 0.375 | 0.586 |

Models that combined VR-elicited expert information with the survey data improved predictive accuracy and produced the most precise parameter estimates. The precision for the parameter estimates in the G model was low relative to the other models (Fig 4), and consistent with its sample size. Adding thermal-imagery data (the GT model) increased the precision and adding information elicited from experts (the G_E model) narrowed the confidence intervals further still (Fig 4). Furthermore, the combined GT_E model generated the most precise estimates of any model for each of the regression parameters.

**Fig 4.**
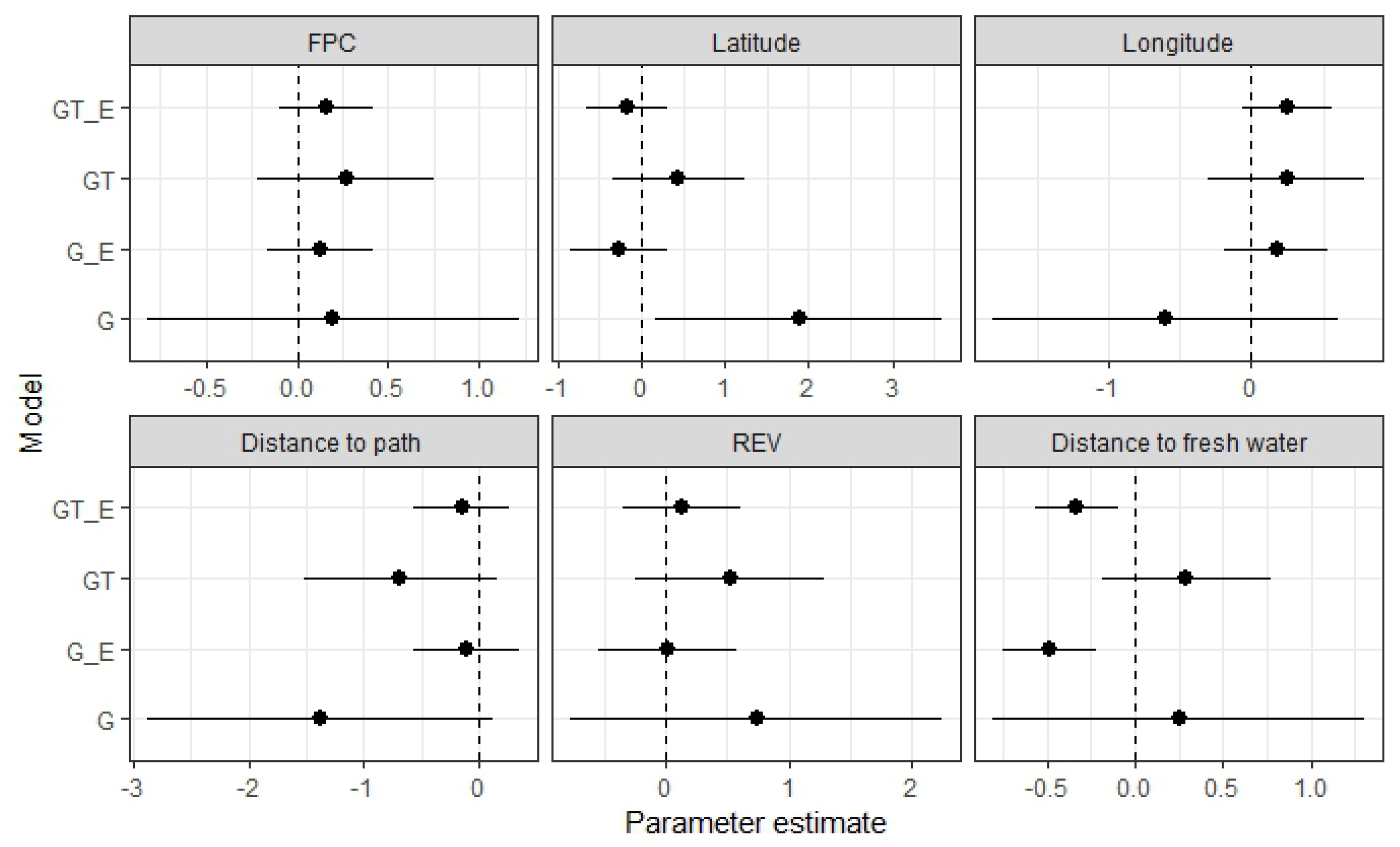
Parameter estimates and confidence intervals. Shown as closed circles with 95% confidence intervals (horizontal bars) for each regression parameter in each model. G = ground survey only, G_E = combined ground-survey and expert-elicitation, GT = ground and thermal-imagery surveys, GT_E = combined ground and thermal-imagery surveys and expert-elicitation.

Latitude and distance to fresh water were the only covariates with significant relationships with koala presence and absence. In the G model, latitude had a significant and positive effect on koala presence/absence, whereas distance to fresh water had significant and negative effects in the G_E and GT_E models (Fig 4). For this latter covariate, there was also a change in the mean direction of its effect among the models, having positive (albeit non-significant) effects in the G and GT models and negative (significant) effects in the G_E and GT_E models (Fig 4).

The predicted presence/absence of koalas in the study area also differed among models (Fig 5). Observer bias was apparent in the G-model predictions, for which presence of koalas was strongly predicted in the northern, more open parts of the study area near residential housing and away from the river, where access would be easy during ground surveys (cf. Figs 1 and 5). In contrast, the GT model predicted that koalas presence would be less likely in the open areas, and more likely closer to the bend of the Logan River where most thermal hot-spots had been identified (Figs 1 and 5). Predictions of koala presence were even less likely in the northern and open areas of the park when the expert information was included in the models (i.e. G_E and GT_E), with the greatest probability of presence occurring in the southwest corner of the park close to the river (Fig 5).

**Fig 5.**
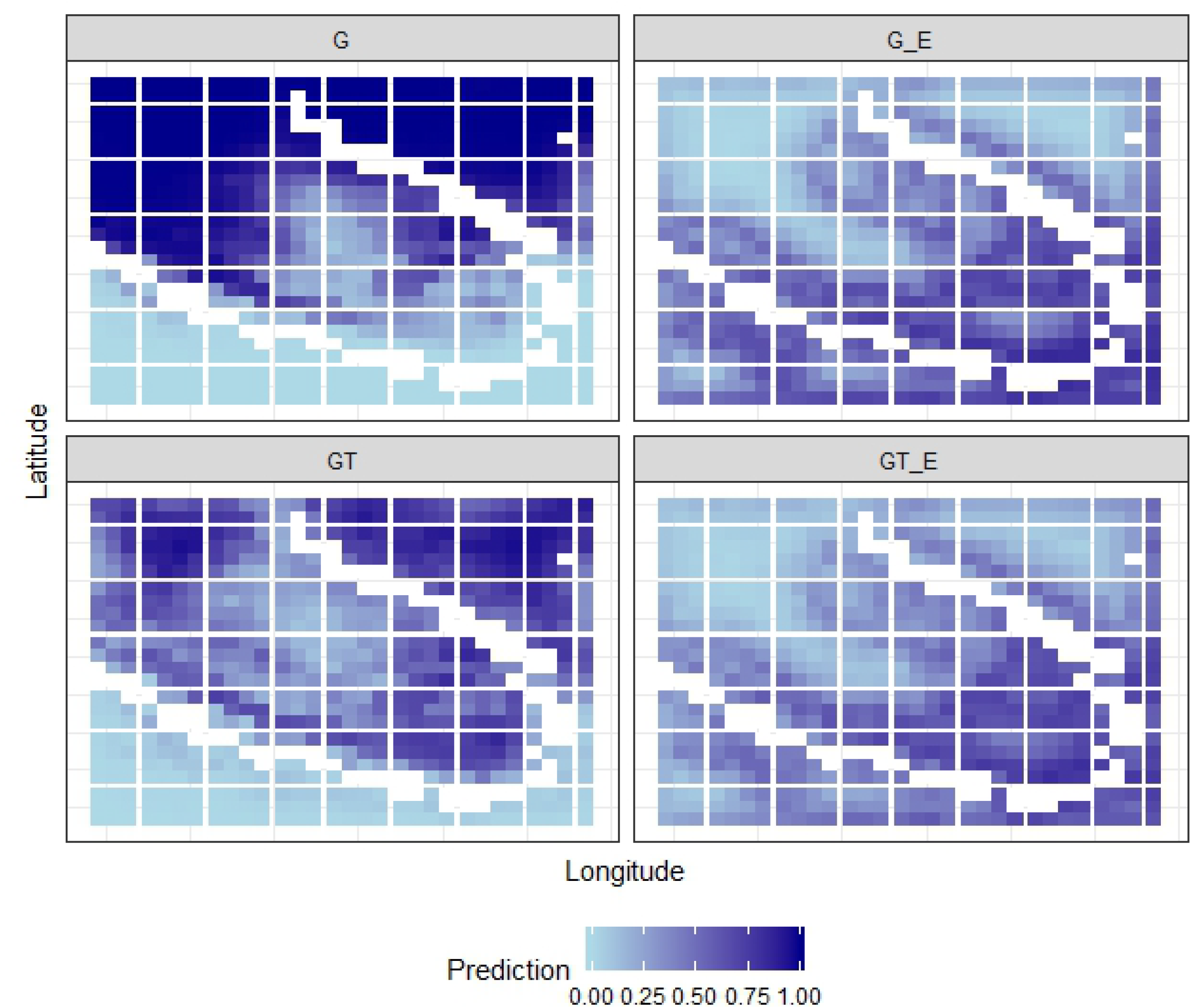
Model predictions. Probability of koala presence/absence in the study area as predicted by each model, shown on a continuous scale of 0 (absence, light blue) to 1 (presence, dark blue), with the Logan River shown in white. G = ground survey only, GT = ground and thermal-imagery surveys, G_E = combined ground-survey and expert-elicitation, GT_E = combined ground and thermal-imagery surveys and expert-elicitation.

## Discussion

Like many cryptic species, koalas are notoriously difficult to detect in the field [15,40]. That they are equally likely to be found in habitat of presumed low or high quality exacerbates their imperfect detection, and observation data are thus highly prone to false negatives [41,42]. Species distribution models typically perform poorly when there is imperfect detection; even models using covariates selected on the basis of the best available knowledge will still perform poorly when misclassifications are present [43]. Results from our case study show how emerging technologies can be harnessed to improve observation-based distribution models for vulnerable and protected species, particularly for cryptic ones like the koala.

The emerging technologies helped to improve models by reducing survey bias [44], specifically by (i) increasing sample size, (ii) sampling in areas that were hard to access, and (iii) generating absences with a high level of confidence such that logistic regression could be used rather than models for presence-only or pseudo-absence data [19]. We also minimized cognitive bias and thus potential under- or overestimation of probabilities during the expert elicitation process [31,45] by using a simple elicitation design (i.e. asking one question about habitat suitability and the uncertainty in that answer only) and randomizing the order in which images were shown. Survey bias was strongly apparent in the ground-survey only model, with koala presences predicted in open areas near residential housing and places that were easily accessible by observers; false negatives that were corrected upon inclusion of the RPAS and/or expert data (Fig 5). Consequently, classification accuracy of the model improved as did the ability to predict koala presences at unobserved locations. This is important because accurate species distribution maps provide information critical for conservation [46], especially given that anthropogenic impacts related to habitat fragmentation and climate change continue to alter species abundances and ranges [2,47]. Improvements in accuracy and, in particular, reductions in false negatives, will provide the information needed to ensure surveillance, habitat restoration and protection measures are implemented in areas most likely to yield positive conservation outcomes [48].

Models combining ground-based observations with data from thermal-imagery and VR-elicited expert information had the most precise regression parameter estimates (Fig 4). However, few covariates had significant relationships with the presence and absence of koalas in the study region. This was the case even when the data from experts and thermal imagery were included in models and despite canopy cover (related to FPC), proximity to roads and dogs (for which distance to paths was proxy) and the presence of food-tree species (*Eucalyptus* spp.) all having been identified previously as important [16,49,50,51]. The relatively coarse scale of the covariates generated from GIS data and their limited range of values within the small study area (S1 Data) may have contributed to these ‘negative’ findings. However, when additional information from experts was included in the models fit to ground-based and/or thermal data, there was a significant relationship between koala presence/absence and distance to fresh water, with greater probability of koalas being present in areas closer to the river. The leaves koalas eat provide most of their water needs, so this covariate may act as a surrogate for tree condition and leaf moisture content [52,53]. Areas surrounding prominent sources of fresh water in other parts of southeast Queensland, for example North Stradbroke Island, also support relatively high numbers of koalas [54]. Such findings thus provide support for research that suggests extreme events, such as droughts and heatwaves, that affect water availability will have both direct and indirect consequences for koalas and other endothermic species that use evaporative cooling for thermoregulation [55,56].

While our study is relatively small in size, both in terms of its spatial extent and the number of koalas observed from the ground, it provides a proof-of-concept that the certainty of models can be increased by including additional information from new technologies and expert elicitation in combination with traditional statistical modelling. Specifically, we demonstrated how ground- and/or RPAS-based observation data can be combined with VR expert-elicitation data to provide a more comprehensive, precise and accurate species distribution model. In the current era of accelerated, human-induced biodiversity loss and high extinction risk [2,57], we need to be innovative and creative about how we capture data and generate information to characterize biodiversity variables for timely and effective conservation. Otherwise, limited knowledge about rare, cryptic and of-concern species, such as differences in habitat needs at different life stages [58], when, where and why organisms move [59], and/or biotic interactions as species ranges shift due to climate change and other anthropogenic impacts [1,57] will continue to constrain conservation efforts. Thus, we do not advocate for an end to ground surveys; ground-collected observational data and existing ecological records will remain vital to ground-truth and combine with other forms of presence/absence data to generate much needed information for conservation [57,60]. Furthermore, the methods we present can be expanded to other wildlife species or places where little to no data are available, particularly where thermal imaging via RPAS is an appropriate solution to any detection obstacles such as site inaccessibility and habitat complexity (e.g. [61]). Such efforts will help to close data gaps and provide the scientific information needed for enhanced conservation of vulnerable and protected species.

## Data accessibility

The R script and associated data to run the models and produce the results and figures herein are available in the Supplementary Information (S5 R Code and input files).

## Acknowledgements

This research was supported by the Australian Research Centre for Mathematical and Statistical Frontiers (ACEMS) and Queensland University of Technology (QUT). We thank Logan City Council for assistance with organizing ground- and thermal-imagery data collection, and the IFE Research Engineering Facility for operating the Remote Piloted Aerial System (RPAS). The expert elicitation protocol was approved by the QUT Research Ethics and Integrity Committee (# 1600000830) and conducted in accordance with the National Statement on Ethical Conduct in Human Research (2014), Australian Code for the Responsible Conduct of Research (2007) and QUT’s Research Governance Framework. Informed consent was obtained from all participants.

## Supporting information captions

**S1 Data. Koala and habitat data.** Koala presence/absence and habitat data collected during field surveys of the study area or derived from freely available geographic information system (GIS) datasets.

**S2 Protocol. Interview protocol.** Standardized expert-elicitation protocol for practice interviews and expert elicitation.

**S3 Fixed effects tables.** Parameter estimates for the fixed effects in the ground survey only (G), ground and thermal-imagery survey (GT), combined ground-survey and expert-elicitation (G_E), and combined ground and thermal-imagery survey and expert-elicitation (GT_E) models.

**S4 Data. Expert elicitation data.** Virtual-reality elicited information from experts viewing 360-degree images of potential koala habitat.

**S5 R Code and input files.** Script and data files needed to run the observation-only, expert-elicitation and combined models in R statistical software.

